# Studying Effects of PDA Media Strengths on the Growth of *Colletotrichum sublineola* Using MPLEx-Based Integrative Proteomics and Metabolomics Analyses

**DOI:** 10.64898/2026.05.15.724728

**Authors:** Pranav Dawar, Dora Farago, Kevin J. Zemaitis, Audrey Thomas, Priscila M. Lalli, Chaevien S. Clendinen, Vanessa L. Paurus, Theresa F. Law, Erin L. Bredeweg, James M. Fulcher, Jeffery L. Dangl, Qun Liu, Ljiljana Paša-Tolić

**Author notes:** corresponding author -Ljiljana Paša-Tolić.

## Abstract

*Colletotrichum sublineola* (*Cs*), the hemibiotrophic fungus that causes sorghum anthracnose, impacts sorghum grain and biomass crop production worldwide. Although nutrient availability is known to influence development in filamentous fungi, including Colletotrichum species, how *in vitro* nutrient limitation reprograms the *Cs* cellular state remains unclear. We cultured *Cs* on full-strength, half-strength, and one-tenth-strength potato dextrose agar (PDA) to define responses across a nutrient gradient. Nutrient limitation induced a pronounced high-sporulation phenotype, with one-tenth-strength PDA producing the strongest conidiation response, followed by half-strength PDA. To study the underlying molecular programs in each condition, we employed a multiplexed <u>m</u>etabolite, <u>p</u>rotein, and lipid <u>ex</u>traction (MPLEx) protocol for global proteomics and metabolomics. Global proteomics resulted in 4,590 protein identifications, including 204 unique to one-tenth-strength PDA. Among them are proteins linked to sporulation, vesicular transport, glycosylphosphatidylinositol (GPI)-anchor biosynthesis, and <u>c</u>ommon in fungal <u>e</u>xtracellular <u>m</u>embrane (CFEM)-domain proteins. Differential abundance and pathway analyses revealed a broad reduction of central carbon and energy metabolism, including glycolysis/gluconeogenesis, pentose phosphate, pyruvate metabolism, and glyoxylate pathways, together with increased ribosome-related processes, cAMP signaling, and cell-surface remodeling in one-tenth-strength PDA conditions. In addition, correlative metabolomics supported selective metabolic depletion and resource reallocation toward stress adaptation, membrane remodeling, and conidiation, supporting proteomics findings. Together, these data support a starvation-adapted *Cs* developmental state associated with enhanced sporulation, cellular pathway reprogramming, and potential virulence linked preparedness under nutrient-limited growth conditions *in vitro*.

**Graphical Abstract:** 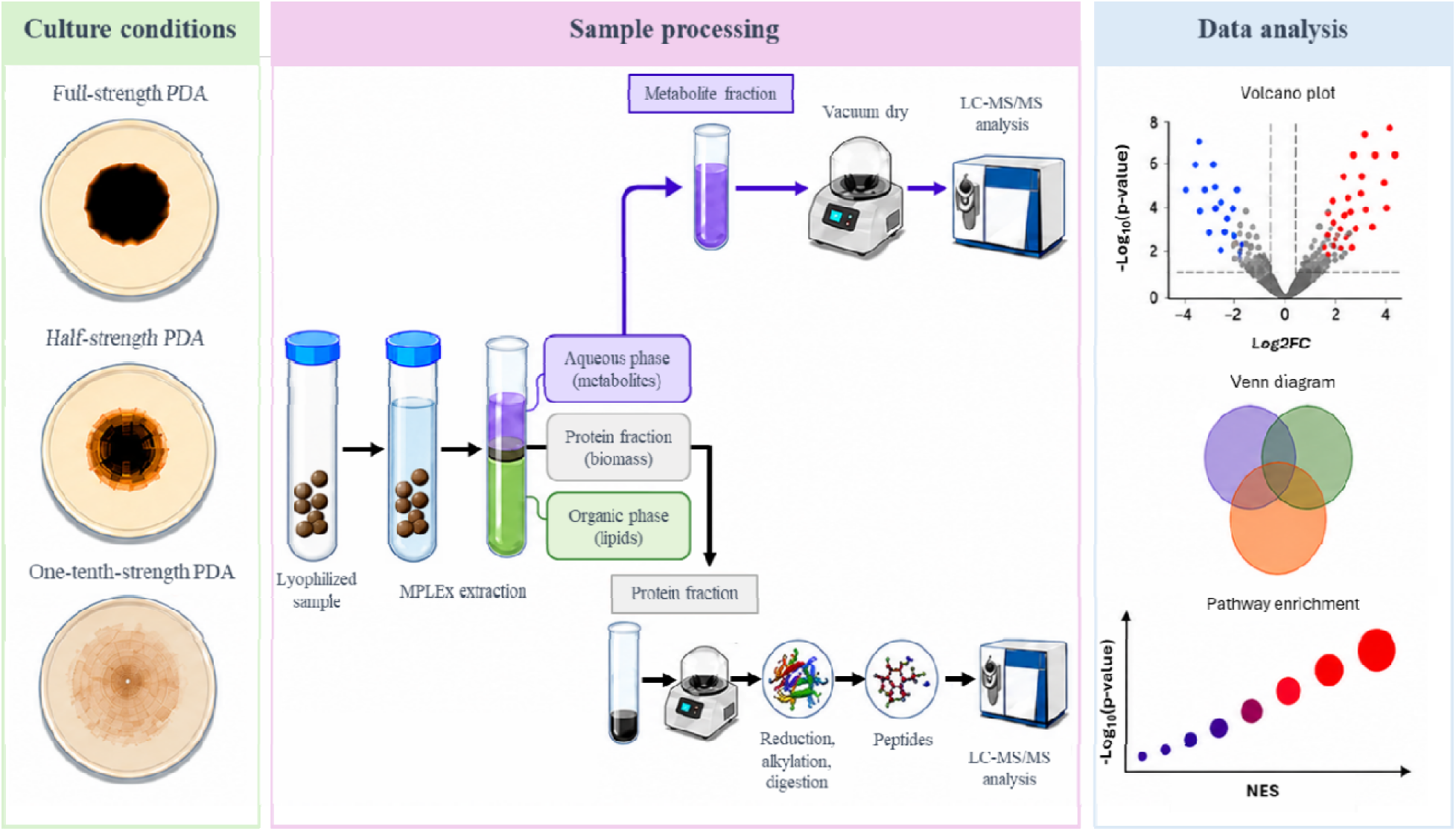

## Introduction

*Colletotrichum sublineola* (*Cs*) is a hemibiotrophic fungal pathogen that causes sorghum anthracnose, one of the most devastating diseases of sorghum worldwide. The genus Colletotrichum can infect multiple tissue types in a wide variety of crop plants and can cause substantial losses in both grain and biomass production (1–4). Because sorghum is a major food, feed, and bioenergy crop, understanding the biology of *Cs* is essential for improving disease management and crop resilience. A major but relatively underexplored factor in *Cs* pathogenicity is nutrient availability. For most filamentous fungal species, nutrient stress along with other environmental stimuli can act as a developmental signal that reshapes fungal physiology before host infection (5). For example, in related *Colletotrichum* systems, before host penetration, spores must rely on internal reserves to support germination/appressorium formation, and early invasive growth under nutrient-poor conditions (6–12). Additionally, nitrogen deprivation, in particular, has been linked to metabolic reprogramming, synchronous pre-infection development, appressorium differentiation, and virulence-associated functions, while regulators of nitrogen metabolism and nutrient transport systems can strongly influence vegetative growth, conidiation, infection related development, and pathogenicity (6, 9, 10, 12). These observations suggest that nutrient limitation may play a role in reprograming *Cs* for infection-related development rather than simply suppressing growth.

In *Cs*, most molecular studies to date have focused on host resistance, comparative genomics, and transcriptomics during the sorghum-pathogen interactions (13, 14) (15, 16). However, the effect of nutrient limitation stress on the growth and development of *Cs*, especially at the combined protein and metabolite levels, remains much less resolved. This represents an important gap to characterize how nutrient limitation reshapes the molecular state of pre-infection *Cs*, for example sporulation capacity, stress adaptation, and virulence associated preparedness, remains largely unknown. To address this gap, we utilized full-strength, half-strength, and one-tenth-strength potato dextrose agar (PDA) to create a controlled nutrient gradient while keeping the base medium constant. Full-strength PDA represents a nutrient-replete condition, half-strength PDA captures an intermediate state, and one-tenth-strength PDA imposes nutrient limitation. It is well known that global as well as low-input proteomics and metabolomics approaches (17–19) successfully capture the cellular stress-responses at functional (enzymes, transporters, stress-response proteins, secreted factors, and developmental regulators) as well as biochemical (storage reserve mobilization, redox balance, osmotic adjustment, and secondary metabolite production) levels, respectively. By comparing *Cs* proteomic and metabolomic responses across these three conditions, we aim to explore how nutrient availability influences fungal growth strategies, including biomass accumulation, sporulation, starvation adaptation, and virulence-related developmental programming, in the absence of host tissue. Because studies involving more than one ‘omics measurement modality (20–22) have been shown to capture a more “complete” outlook at a systems biology level, we applied MPLEx-based (17) global proteomics and metabolomics to Cs samples to reveal the molecular networks associated with nutrient limitation stress.

## Results

### Nutrient Depletion Drives a High-Sporulation Phenotype and Distinct Condition-Specific Proteins in *Cs*

Nutrient availability is a major determinant of *Cs* developmental decisions, and nutrient depletion, including carbon, and nitrogen, is often sufficient to redirect resources from vegetative growth toward asexual reproduction as well as virulence (5, 8–10, 12, 23). To examine whether starvation-like nutrient conditions promote sporulation, we cultured *Cs* across different strengths of PDA concentrations, namely full-strength PDA (**Figure 1A)**, half-strength PDA (**Figure 1B)**, and one-tenth-strength PDA (24) (**Figure 1C)**, with full-strength PDA being the baseline medium as it is a widely used as a reproducible platform for culturing and maintaining stable colony phenotype of many filamentous fungi and reliably supports vegetative growth in *Cs* (25–27). As expected, we observed a pronounced shift toward a high-sporulation state, in response to nutrient-depletion, and this was accompanied by distinct growth phenotypes relative to full-strength PDA (**Figure 1D)**. We observed a significantly high production of conidia under one-tenth-strength (p-value of 4.748e-10) and half-strength (p-value of 0.002565) PDA conditions relative to full-strength PDA conditions (**Supplementary Data S1**), with the highest mean number of conidia produced at one-tenth-strength PDA conditions, indicating that the nutritional stress imposed in this study is sufficient to drive *Cs* toward robust asexual reproduction. In addition, *Cs* exhibited reduced vegetative growth and increased radial growth under nutrient-limited conditions.

**Figure 1.**
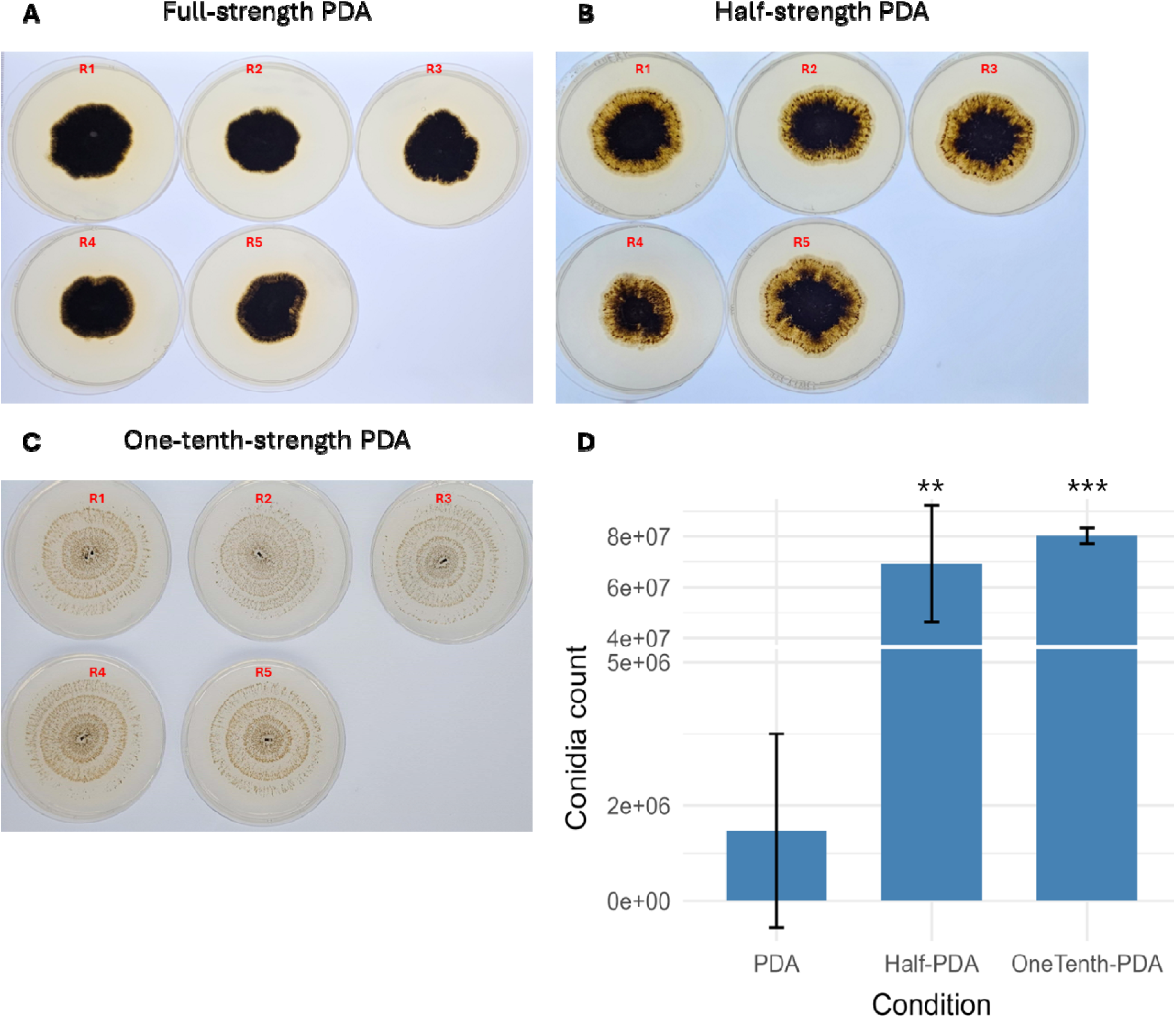
Phenotype of 2-week-old *Cs* cultures. grown on **(A)** full-strength PDA, **(B)** half-strength PDA, and **(C)** onetenth- strength PDA media. **(D)** Bar plot showing the mean number of conidia extracted from 2-week-old cultures under each medium condition. * represents the significance level based on unpaired Welch’s two-tailed t-test , where for full-strength vs one-tenth-strength PDA (*** pvalue of 4.748e-10), and for full-strength vs half-strength (** pvalue of 0.002565) PDA.

To link these observations to proteome-level remodeling, we performed label-free bottom-up proteomics across biological replicates for each condition and QC-verified high-quality identifications by using conservative criteria (≥3 quantified observations per condition across five biological replicates). In total, we identified 53,862 unique peptides and 4,590 proteins across all conditions. On average, we quantified 4,013 ± 57 (mean ± s.d.) proteins in half-strength PDA, 3,379 ± 108 in full-strength PDA, and 2,783 ± 134 in one-tenth-strength PDA, benchmarking the deep proteomics coverage in our datasets (**Figure 2A**). Biological replicates were highly reproducible, with mean Pearson correlations ranging from 70 to 92% within each condition (**Supplementary Figure 1**). Consistent with this, median coefficients of variation (CVs) were 0.264, 0.221, and 0.287 for full-strength, half-strength, and one-tenth-strength PDA conditions, respectively (**Figure 2B**). Additionally, principal component analysis (PCA) indicated that the PC1 pseudo-dimension correlated significantly with growth condition (p-value < 0.01) and accounted for ∼21% of the observed variation, allowing for clear separation of one-tenth strength PDA condition from full-strength and half-strength PDA conditions (**Figure 2C**). Together, these results indicate that our proteomics dataset robustly captures the biological variation induced by nutrient conditions.

**Figure 2.**
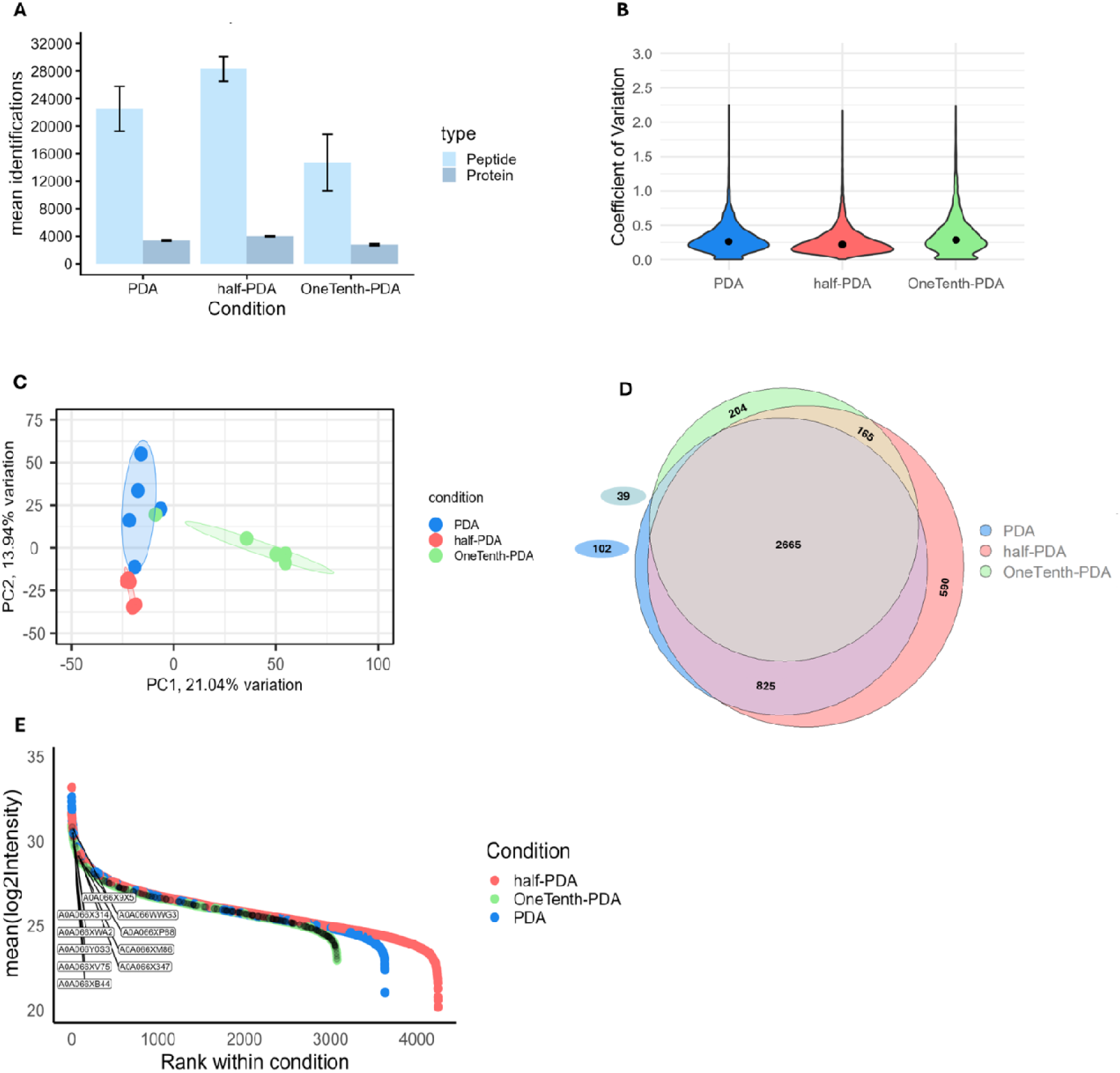
Proteomic profiling of *Cs* grown on full-strength, half-strength, and one-tenth-strength PDA media. **(A)** Mean peptide and protein identifications prior to imputation across the three growth conditions (n = 5). **(B)** Protein-level (n = 4,590) coefficients of variation for *Cs* samples under each medium condition. **(C)** Proteomics informed principal component analysis (PCA) of *Cs* samples grown on full-strength, half-strength, and one-tenth-strength PDA. **(D)** Venn diagram showing shared and unique proteins identified across growth conditions, considering proteins detected in more than two replicates (n ≥3). **(E)** Log2-transformed mean protein intensities for proteins observed in each growth condition with n ≥3. Black points represent the top 10 proteins with the highest intensity from the set of 204 proteins specific to the one-tenth-strength PDA condition.

Most of the proteins identified were shared across the three conditions; however, 204 proteins were uniquely identified in one-tenth-strength PDA (**Figure 2D, Supplementary Data S2**). Upon further investigation, we found several proteins known to be directly and indirectly involved in the sporulation and virulence-related functions in filamentous fungi, namely, A0A066X743 (SPO7-like) (28–30), A0A066X6N6 (Conidiophore development protein hymA) (31, 32), glycosylphosphatidylinositol (GPI)-anchor biosynthesis pathway (A0A1B7YF74, A0A066XBY7, A0A066XE37) (33–36) and SNARE interactions in the vesicular transport pathway (A0A066XFU6, A0A066XJQ8) (37–39). Moreover, the one tenth strength PDA-specific proteins exhibited a broad intensity distribution, compared to all the proteins identified under the one-tenth-strength condition (**Figure 2E**). Among the 204 uniquely-detected proteins, iron/host interface (CFEM) (40, 41) domain-containing proteins were especially abundant, ranking within the top 10th percentile when proteins were ordered from highest to lowest abundance based on log2 mean intensity across biological replicates. In addition, ∼13% of the proteins unique to one-tenth-strength PDA were annotated as “uncharacterized”, underscoring a need for complete functional annotation of the *Cs* proteome. For example, one of the uncharacterized protein (A0A066XP88), within the top 10th percentile, is found to contain a COX7a domain (42), which is associated with stabilization of the cytochrome c oxidase complex in mitochondrial respiration. Expression of this protein uniquely under nutrient limitation points to a potential connection between mitochondrial respiratory function and a role in nutrient-stress responses or sporulation.

### Proteomics-informed *Cs* pathways are involved in nutrient depletion stress response

Having identified sporulation-specific proteins unique to one-tenth-strength PDA condition samples, we wanted to understand the overall *Cs* response to nutrient limitation stress relative to full-strength and half-strength PDA conditions. To this end, we performed differential abundance analyses on the proteome captured across all three experimental conditions. Using a log2 fold-change cutoff of ±1 and an adjusted p-value threshold of 0.05, we identified a total of 740 unique differentially abundant proteins (DAPs) associated with nutrient limitation stress (**Figure 3A**). 520 DAPs were detected when one-tenth-strength PDA samples were compared with full-strength PDA, and 461 DAPs were detected when one-tenth-strength PDA samples were compared with half-strength PDA (**Supplementary Data S3, S4**). On the other hand, only 71 DAPs were identified when half-strength PDA samples were compared with full-strength PDA (**Supplementary Data S5**). KEGG-based gene set enrichment analysis (GSEA) of these 71 DAPs did not reveal any significantly enriched pathways, suggesting that half-strength PDA remains largely comparable to full-strength PDA at the pathway level, and likely does not impose a strong nutrient limitation stress response.

**Figure 3.**
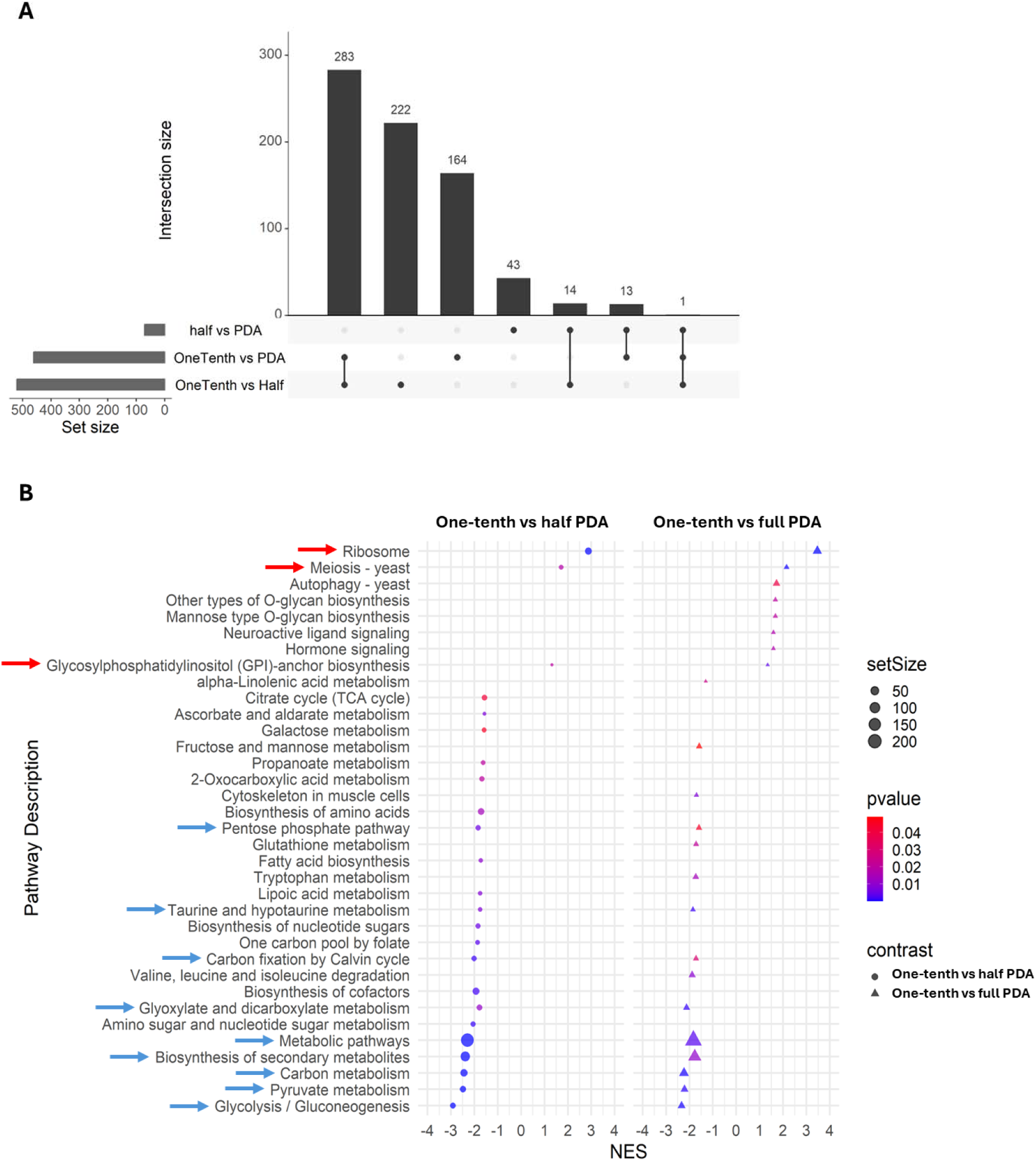
Differential proteomic responses of Cs grown on varying PDA strengths. (A) UpSet plot showing unique and overlapping differentially abundant proteins (DAPs) identified in the comparisons of half-strength vs full-strength, one-tenth-vs full-strength, and one-tenth-strength vs half-strength PDA conditions, using thresholds of |log2FC| ≥ 1 and BH-adjusted p < 0.05. (B) Gene set enrichment analysis using the KEGG pathway database, performed on ranked DAPs, showing positively and negatively enriched pathways for each comparison, shown by normalized enrichment score (NES) values. Red arrows represent the positively-enriched and blue arrows represent negatively enriched pathways shared across both comparisons.

Downstream analysis of DAPs from the comparisons involving one-tenth-strength PDA revealed that 283 DAPs were shared between the two contrasts, while 222 and 164 DAPs were unique to the one-tenth-strength versus half-strength and one-tenth-strength versus full-strength comparisons, respectively (**Figure 3A**). Note, all shared DAPs exhibited similar abundance trends across both contrasts, indicating that nutrient limitation draws out a consistent proteomic reprogramming within this shared protein subset, independent of whether the reference condition is full-strength or half-strength PDA. Under nutrient limitation stress, pathways associated with primary and secondary metabolism, such as glycolysis/gluconeogenesis (chig00010), the pentose phosphate pathway (chig00030), pyruvate metabolism (chig00620), and glyoxylate/dicarboxylate metabolism (chig00630), as well as secondary metabolism pathways (chig01110), were significantly less enriched (**Figure 3B**). In contrast, proteins associated with ribosome-related (translation and protein turnover; chig03010), cell cycle-linked (cAMP signaling; chig04113) and virulence-related (GPI anchor biosynthesis; chig00563) pathways were consistently over-enriched across nutrient-limited conditions (**Figure 3B**). Together, these observations point to extensive pathway shifts in *Cs* under nutrient limitation, characterized by reduced abundance of proteins involved in core metabolic pathways and increased investment in proteostasis, intracellular signaling, and cell-surface remodeling. Such changes are consistent with a shift away from energy-intensive metabolic pathways and toward cellular reprogramming associated with stress adaptation and developmental transitions, including sporulation, as found previously (34, 35, 43–45).

### Protein covariation analysis highlights *Cs* proteins with putative roles in sporulation and virulence

We next moved beyond differential abundance analyses and explored our proteomics data to examine protein covariation under nutrient limitation stress. Because covariation patterns can reflect shared function, co-regulation, and biophysical relationships (46), we asked whether protein covariation patterns in our data could provide insight into putative protein-protein interactions and help identify proteins that act together to regulate stress-responsive pathways (“guilt-by-association”). To enable robust network inference, we used previously identified 4,590 QC-qualified proteins (≥3 quantified observations per condition across five biological replicates), imputed the missing values using k-nearest neighbors (KNN), and then applied Gaussian mixture model (GMM) clustering (k = 27) to identify groups of high confidence covarying proteins using uncertainty cut-off of ≤0.1. Further to visualize the resulting consensus network, we computed all pairwise Pearson correlation coefficients and summarized these relationships at the cluster level (**Figure 4A**). In order to identify potential proteins involved in sporulation- and virulence-related pathways, we specifically used proteins (A0A066Y192, A0A066XJE7, A0A066XBF5, A0A066XBE1, A0A066XT04, A0A066XBI2) mapped to cell-cycle related pathway (chig04113) and GPI-anchor biosynthesis (chig00563; A0A066X3R0) pathways (**Figure 3B**) as bait proteins. Notably, these proteins were mapped to clusters 8, 13, and 18 of the covariation networks, which contain 226, 106, and 141 additional proteins, respectively. At the cluster level, clusters 8 showed strong positive mean correlation with cluster 13 (r = 0.83) and cluster 18 (r = 0.84), whereas clusters 13 and 18 showed no mean correlation (**Figure 4A**). This pattern suggests that cluster 8 proteins share coordinated abundance behavior with proteins mapped to both cluster 13 and 18, whereas clusters 13 and 18 likely represent distinct modules.

**Figure 4.**
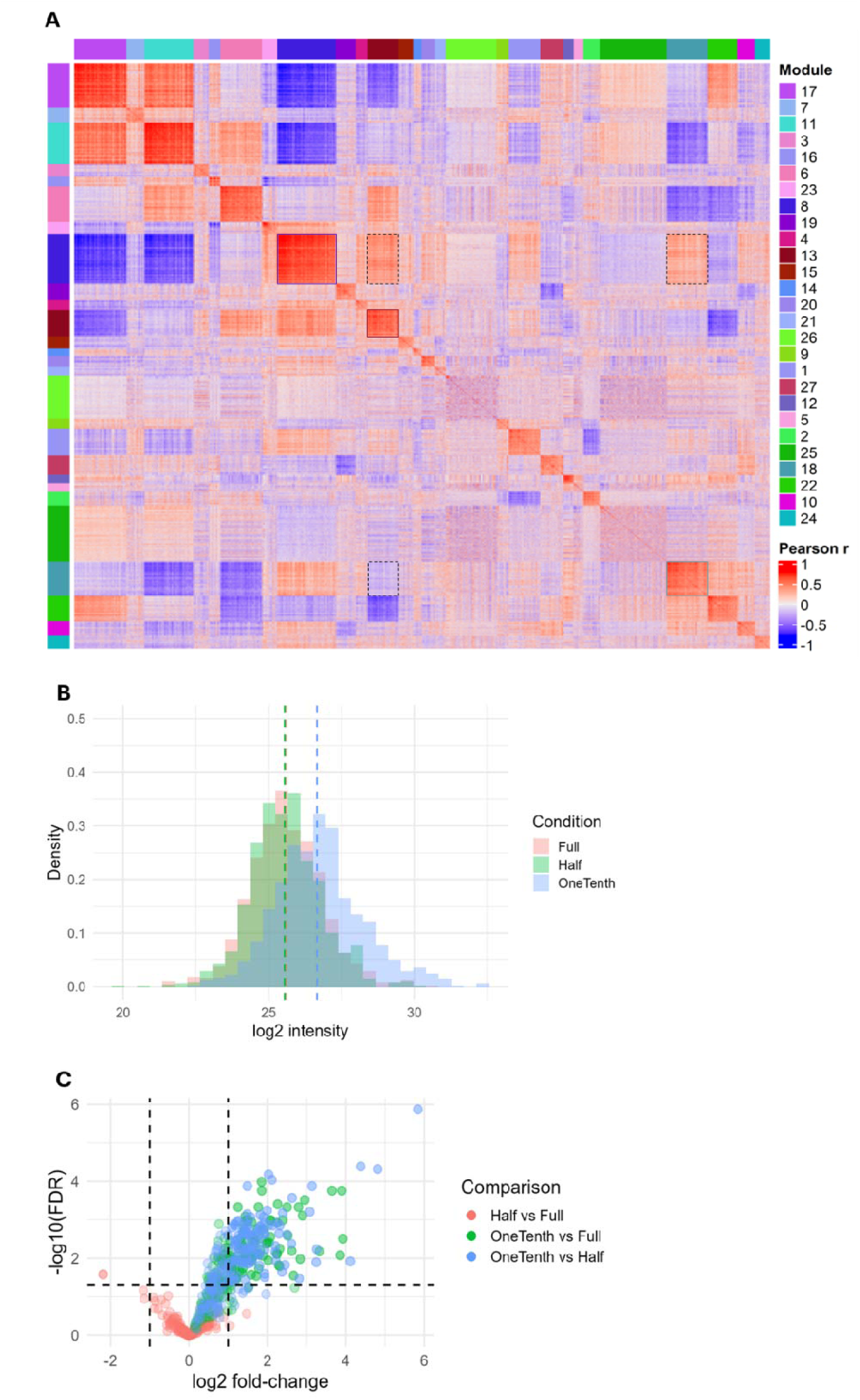
Gaussian mixture model-based clustering reveals coordinated protein abundance patterns in Cs. (A) GMM clustering of all imputed proteins using a pairwise Pearson correlation matrix identified 27 clusters, uncovering protein co-variation networks and highlighting clusters 8, 13, and 18 in association with the previously identified chig04113 and chig00563 KEGG pathways. (B, C) Proteins assigned to cluster 8 exhibited a consistent increase in abundance in the one-tenth-strength PDA condition.

To assess whether proteins assigned to clusters 8, 13, and 18 were associated with nutrient limitation stress responses, we examined their abundance patterns across all growth conditions to determine if any trends were evident. Interestingly, proteins in cluster 8 displayed a consistent increased abundance trend in one-tenth-strength PDA (**Figures 4B, C**), whereas no clear trend was observed for proteins mapped to clusters 18 and 13 (**Supplementary Figure 2A-D**). 116 of the 227 (∼51%) proteins mapped to cluster 8 showed significant increased abundance, with log2FC ≥ |1| and adjusted p < 0.05 (**Figure 4C**).

Additionally, **Table 1** highlights selected cluster 8 proteins that have been previously identified to play a role in sporulation and/or virulence in filamentous fungi but have not been exclusively explored in *Cs* (complete list of cluster 8 proteins is shown in **Supplementary Data S6**). Note, not all the proteins listed in **Table 1** were captured as DAPs, reiterating the significance of protein-protein covariation analysis (**Supplementary Data S6**). Collectively, these results suggest that proteins mapped to cluster 8 are strongly associated with the nutrient limitation stress response and may represent a coordinated module potentially linked to sporulation- and virulence-related pathways, making them the candidate targets for developing biocontrol strategies.

**Table 1.** Select cluster 8 proteins implicated in sporulation and/or virulence in *Cs*.

| **Protein ID** | **Annotation** | **Biological function** | **Reference** |
| --- | --- | --- | --- |
| A0A066Y192;<br>A0A066X0C4 | cAMP-dependent protein<br>kinase; cAMP-dependent<br>protein kinase regulatory<br>subunit | PKA Signaling;<br>conidiation/appressorium<br>development | (45) |
| A0A066X3R0 | Putative peptidase C13 family<br>protein | GPI transamidase complex;<br>cell-wall organization | (33) |
| A0A066WZQ2;<br>A0A066XGV1;<br>A0A066XH75 | Dolichyl-phosphate-mannose-<br>protein mannosyltransferase | Protein O-mannosylation; cell<br>wall/morphogenesis | (47) |
| A0A066X3R1 | Mitogen-activated protein<br>kinase kinase | MAPK signaling; infection<br>development | (48) |
| A0A066XPP0 | Putative calcineurin | Calcium signaling;<br>morphogenesis/stress response | (49) |
| A0A066XXH7 | STT3 oligosaccharyltransferase<br>subunit | N-glycosylation; secretion/cell<br>surface function | (50) |
| A0A066XDW2;<br>A0A066XGL0 | Peroxisomal ATPase PEX1;<br>PEX11 | Peroxisome biogenesis; lipid<br>metabolism | (51) |
| A0A066XEB8 | Sphingolipid C(9)-<br>methyltransferase | Sphingolipid biosynthesis;<br>membrane integrity | (52) |
| A0A066XXR7;<br>A0A066XIB0 | Exocyst complex component<br>SEC5 ; SEC8 | Polarized secretion;<br>hyphal/infection growth | (53) |
| A0A066X4J2 | Outer spore wall protein RRT8 | Spore wall assembly | (54) |

### Metabolomics Validates Proteomic Signatures of Nutrient-Dependent Remodeling

While proteomics revealed broad condition-specific remodeling of the *Cs* cellular machinery under nutrient limitation stress, we reasoned that direct measurement of metabolites would be essential to determine how these protein-level changes corroborate with metabolomic changes. Unlike proteins, which reflect cellular investment in enzymatic and structural capacity, metabolites represent the immediate products and substrates of pathway activity and therefore provide a snapshot of the physiological state of a cell. We therefore performed a global metabolomics analysis from the same sample set to define how nutrient conditions reshape fungal metabolome. Across all samples and ionization modes, we detected 2832 metabolite features, of which 2038 passed QC-filtering criteria outlined in the methods section and were retained for downstream analyses. Similar to the proteomics dataset, biological replicates showed strong within-condition clustering, supporting robust metabolomics analyses. PCA demonstrated clear condition-dependent separation, with one-tenth-strength PDA samples clearly resolving from full-strength and half-strength PDA along PC1, which explained 48.4% of the total variance, whereas PC2 explained 19.7% of the total variance (**Figure 5A**). Additionally, using a cut-off of adjusted p-value < 0.05 and log2FC ≥ |1|, we identified a total of 786, 337, and 353 metabolites significant to half-strength vs full-strength, one-tenth-strength vs full-strength, and one-tenth-strength vs half-strength PDA conditions, respectively (**Supplementary Data S7, S8, S9**). Among these, 386 metabolites were shared across at least two comparisons, indicating nutrient-responsive metabolic remodeling under intermediate nutrient limitation (**Figure 5B**). This observation is particularly significant because the half-strength PDA condition showed comparatively limited DAPs at the proteome level, suggesting that metabolite-level changes may provide a more sensitive readout of an early-stage physiological adaptation to nutrient limitation stress.

**Figure 5.**
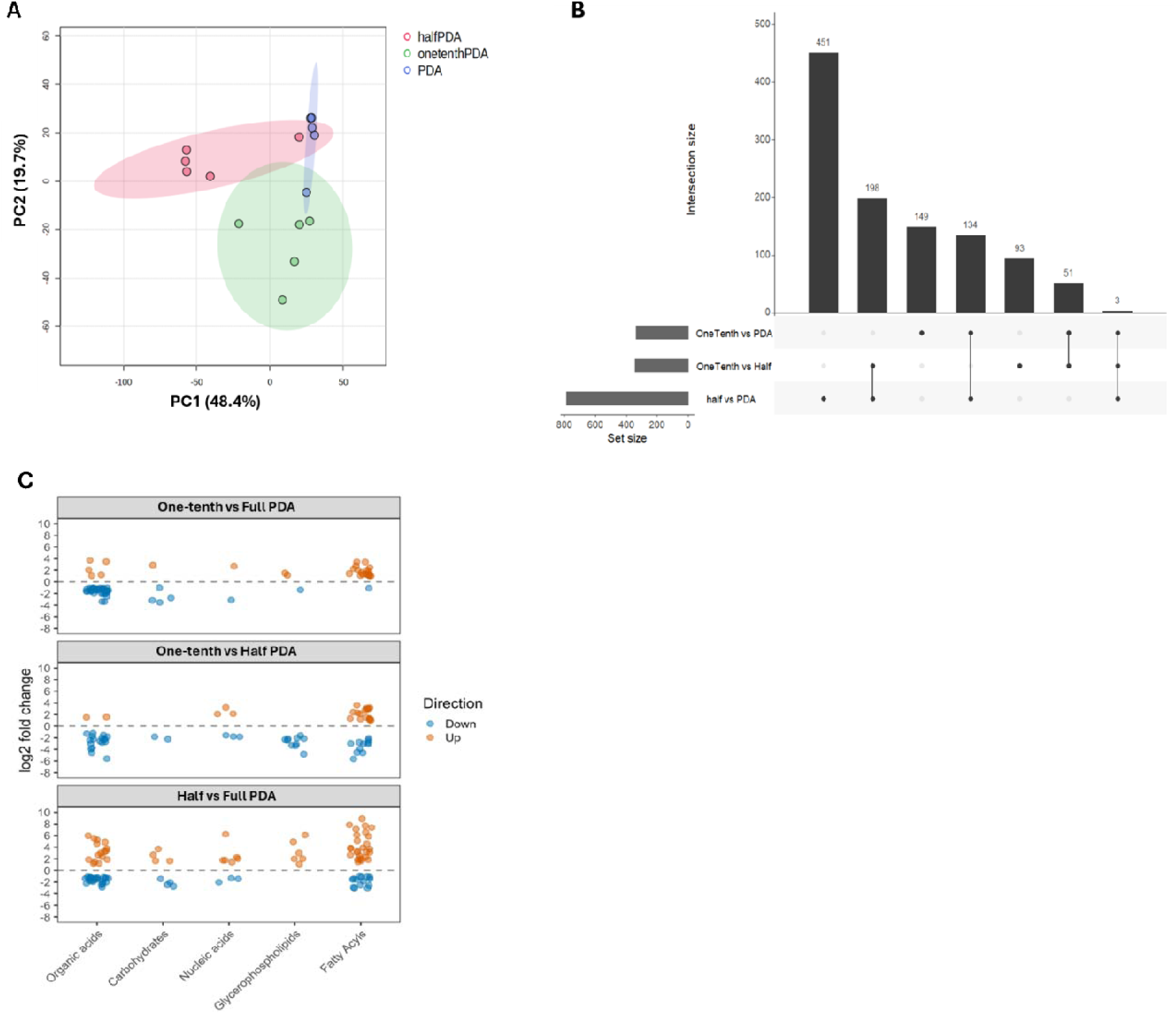
Metabolomic profiling of *Cs* grown on full-strength, half-strength, and one-tenth-strength PDA media. (A) Principal component analysis (PCA) of metabolomics data (n = 2,832) from *Cs* samples grown under full-strength, half-strength, and one-tenth-strength PDA conditions. (B) UpSet plot showing unique and overlapping differentially abundant metabolites (DAMs) identified in the comparisons of half-strength vs full-strength, one-tenth- strength vs full-strength, and one-tenth-strength vs half-strength PDA, using thresholds of |log2FC| ≥ 1 and adjusted p-value < 0.05. (C) Distribution of significantly altered metabolites across major chemical classes, including organic acids, carbohydrates, nucleic acids, glycerophospholipids, and fatty acyls, for all nutrient condition comparisons, using thresholds of |log2FC| ≥ 1 and p-value < 0.05.

Following the metabolite annotation as described in the methods section, we compared the distribution of metabolite fold changes, using p-value < 0.05 and log2FC ≥ |1| cut-offs, across major chemical classes under all nutrient condition comparisons (**Figure 5C**), and found clear class-dependent shifts rather than a uniform metabolic response to nutrient limitation. Among the metabolite classes examined, fatty acyls and organic acids exhibited the largest number of retained features across all contrasts. We found that nutrient limited PDA conditions, in all comparisons, showed a strong and positive accumulation of fatty acyl species. In these data, lipid remodeling was a prominent feature of the nutrient limitation response and remains active across both intermediate and severe nutrient limitation conditions. This is consistent with observations of nutrient limitation in *Aspergillus oryzae,* which has been shown to reshape fatty acid composition and redirect lipid allocation towards stress-associated lipid metabolism (55).

Glycerophospholipids, on the other hand, exhibited condition-dependent remodeling, and showed predominantly negative shifts under one-tenth-strength conditions, but positive shifts in the half-strength versus full-strength PDA comparison. Given glycerophospholipids are central to membrane composition and that their homeostasis is closely linked to morphogenesis, vegetative growth and asexual development, the increased accumulation under half-strength PDA reflects adaptive membrane remodeling compatible with continued vegetative growth as well as conidiation, while decreased abundance under one-tenth-strength PDA is consistent with enhanced phospholipid turnover under severe nutrient limitation (56). Collectively, these lipid metabolite observations support the high-sporulation phenotype observed under one-tenth-strength PDA, as fatty acids and lipid droplets support conidial development, germination, and infection-related morphogenesis in other filamentous fungi (57, 58).

In contrast, organic acids and carbohydrates showed predominantly negative shifts under one-tenth-strength PDA relative to both full-strength and half-strength PDA, indicating depletion of these metabolite pools under nutrient limitation. Because these classes include metabolites associated with central carbon metabolism and readily available biosynthetic intermediates, their reduction is consistent with suppression or rerouting of growth-associated metabolic activity under starvation. Notably, both organic acid and carbohydrate classes displayed broad positive shifts in the half-strength versus full-strength PDA comparison, suggesting that a less stringent nutrient limitation condition does not simply reduce metabolic abundance, but instead induces adaptive metabolite homeostasis. Together, these class-level patterns support a model in which nutrient limitation stress drives progressive metabolic reprogramming in *Cs*, consistent with the proteomics results, which measure a shift away from vegetative, growth-oriented metabolism and toward a starvation-induced state that emphasizes resource reallocation, membrane remodeling, as well as enhanced sporulation.

## Discussion

Results from this study show that nutrient availability strongly shapes the developmental behavior of *Cs*. As demonstrated previously, we also showed that under nutrient limitation, especially under one-tenth-strength PDA, *Cs* produced significantly more conidia (**Figure 1D**), indicating a clear shift away from vegetative growth and toward asexual reproductive state. Half-strength PDA conditions also increased conidiation compared with full-strength PDA, but notably fewer conidia than one-tenth-strength PDA conditions. The enhancement of conidiation suggests that nutrient depletion acts as an important cue for asexual reproduction in *Cs* likely as part of a survival strategy as speculated before (23). It was previously established that sporulation develops contingent on unfavorable conditions because it allows the organism to persist, disperse, and potentially colonize new environments. Our observations are consistent with this broader view, and indicate that *Cs* responds to nutrient limitation stress in a similar way (5, 24, 30).

In order to understand how the sporulation developmental shift is regulated, we performed MPLEx-driven proteomics and metabolomics analyses from same sample. Both proteomics- and metabolomics-informed analyses provide strong support for the phenotypes observed, and showed that nutrient depletion is associated with coordinated remodeling of the *Cs* pathways rather than a simple reduction in biosynthetic activity. Bottom-up proteomics analyses revealed a distinct set of 204 proteins, including factors linked to conidiophore development, vesicular trafficking, GPI-anchor biosynthesis, and CFEM-domain proteins, unique to one-tenth-strength PDA conditions (**Figure 2D**). These observations are notable because they connect nutrient limitation stress not only to sporulation-associated developmental machinery but also to cell-surface and host-interaction processes that are frequently linked to fungal virulence (35, 40, 41). For example, CFEM-domain proteins, found abundant in one-tenth-strength conditions, are known to be associated with host interface functions, iron acquisition, and pathogenicity-related processes in filamentous fungi. Their increased abundance under severe nutrient limitation suggests that starvation in *Cs* triggers a broader adaptive response that couples reproductive output, such as conidiation, with other traits important for survival and host interaction. At the global pathway level, we found that nutrient depletion stress response was associated with reduced abundance of proteins involved in central carbon metabolism and energy-generating pathways, alongside increased abundance of proteins linked to protein turnover, intracellular signaling (cAMP-PKA signaling; **Table 1**), and cell-surface remodeling (GPI anchoring; **Table 1**). The collective downregulation of glycolysis/gluconeogenesis, pentose phosphate pathway, pyruvate metabolism, and glyoxylate/dicarboxylate metabolism (**Figure 3B**) suggests that *Cs* reduces investment in growth-supporting metabolic networks, which is consistent with a physiological shift from biomass accumulation to stress adaptation (**Figures 1A, B, C**). In parallel, increased abundance of ribosome-related pathways during nutrient limitation points to selective protein synthesis and turnover, a mechanism for recycling cellular amino acid reserves essential for the adaptive response to nutrient limitation stress. In addition, cAMP-PKA signaling is a well-established regulator of conidiation, appressorium development, and pathogenicity (43–45), whereas GPI-anchor biosynthesis is central to fungal cell-wall architecture and morphogenesis (34–36). The coordinated regulation of these pathways suggests that nutrient limitation stress does not simply suppress growth, but promotes a broader developmental transition that primes *Cs* for functions that are advantageous not only for reproduction but also for host colonization and environmental adaptation.

Protein-protein covariation analysis extends these findings by identifying coordinated protein modules that participate in this transition (**Figure 4A**). In particular, cluster 8 emerged as a candidate module (**Figures 4B, C**) because it shows a clear abundance increase in one-tenth-strength PDA and also contains many proteins, including a subset that were not identified as differentially abundant but have previously been linked to sporulation, morphogenesis, secretion, cell-wall assembly, membrane biogenesis, and virulence-related pathways in other filamentous fungi. These include components of PKA (45) and MAPK (48) signaling, spore wall assembly, hyphal growth, morphogenesis and stress response (**Table 1**). Because many of these proteins have not been functionally characterized in *Cs*, cluster 8 provides a focused set of candidates for future genetic or biochemical studies aimed at dissecting the regulatory basis of sporulation and their possible roles in defining the pathogenic potential of *Cs*.

Metabolomics data further reinforce the phenotypic observations that nutrient depletion induces a major physiological state change in *Cs*. Interestingly, metabolite-level changes were already evident under half-strength PDA conditions, even though the proteome at half-strength PDA conditions remained similar to full-strength PDA conditions. This suggests that the metabolome provides a more sensitive readout of early adaptation to a less stringent nutrient limitation stress, capturing rapid biochemical adjustments that precede detectable changes in steady-state protein abundance. Class-level metabolite profiles also support proteomics-informed resource reallocation under nutrient limitation. For example, the depletion of organic acids and carbohydrates (**Figure 5C**) under one-tenth-strength PDA is consistent with suppression of pathways associated with carbon cycle, and a general redirection of resources away from growth-associated metabolism. The accumulation of fatty acyl species across all nutrient limitation stress comparisons suggest that lipid remodeling is a central feature of the *Cs* stress response (**Figure 5C**). Because lipid reserves are widely associated with conidial maturation, germination capacity, and infection-related development in filamentous fungi, the observed metabolite dynamics support the high conidiation phenotype observed under nutrient limitation conditions.

It is important to note that this study is the first, to our best knowledge, to study nutrient limitation stress response in *Cs in vitro*. While our data does not directly demonstrate enhanced virulence under nutrient limitation, it suggests that nutrient limitation stress may prime *Cs* not only to promote reproduction, but also to be ready for opportunistic host interaction, necessary for its survival. The measured proteomics and metabolomics response is consistent with the idea that nutrient limitation stress can serve as a cue for developmental states that may improve *Cs* readiness for infection. The molecular details provided here provide targets to be tested for their involvement in virulence and infection in future targeted approaches. This study highlights important gaps in current *Cs* reference annotation as a substantial fraction of proteins unique to one-tenth-strength PDA were uncharacterized, including highly abundant candidates such as A0A066XP88, containing a COX7a-related domain, which may link mitochondrial respiratory processes to starvation adaptation (42). Such gaps represent an important but underexplored layer of *Cs* response. Further functional annotation is needed to identify key determinants of *Cs* development that remain hidden within incompletely annotated regions of the proteome.

## Material and methods

### Growth and culture conditions

The sorghum-specific *C. sublineola* pathotype *FSP237* were originally isolated from field sites in Texas and were propagated from single-spore isolate (59, 60). *FSP237* was selected for this study based on its highly aggressive disease phenotype within eighteen sorghum differential lines (60). Cs cultures were grown from frozen Whatman paper covered in dormant fungal spores kept in -80 C. To revive the stock, those frozen chunks of agar were then placed in the middle of a petri dish with full-strength PDA (Potato Dextrose Agar, Millipore Sigma, cat. no. 70139) and incubated at 28 C. Subcultures were generated by cutting 1-2 mm mycelial plugs from the actively growing edge of a 7 day old colony on full-strength PDA (Potato Dextrose Agar, Millipore Sigma, cat. no. 70139). The plugs were transferred onto full-strength, half-strength, and one-tenth-strength PDA (MP Biomaterials, cat no. 1008517) in 9 cm Petri dishes with a final percentage of agar (Millipore Sigma, cat. no. A1296) at 1.5%, with five replicate plates per condition. All cultures were incubated at 28 °C in the dark for 14 days. On day 14, each culture was flooded with distilled water, and the resulting mycelial biomass was gently scraped to avoid disturbing or contaminating the underlying medium. The biomass was transferred to 2 mL microcentrifuge tubes, resuspended, and washed twice with 18.2 Ω Milli-Q water to remove any residual media. The cleaned biomass pellet was then frozen at -80 °C and lyophilized overnight. Following lyophilization, equal amounts of fungal biomass were aliquoted into 1.5 mL microcentrifuge tubes to standardize sample input across all conditions.

### Conidia/spore counting

Conidia were collected from above-mentioned separate PDA-strength cultures, by washing the colony surface with 5 mL of 18.2 Ω Milli-Q water. The suspension was filtered through 70 µm pore size filter to remove mycelial debris. Spore concentration was estimated using a Bright-Line hemocytometer. Briefly, 10 µL of a thoroughly mixed conidial suspension was loaded onto the hemocytometer, and conidia in the 5 major squares were counted using a light microscope in 2 technical replicates. Total spore count was calculated as the average number of conidia per major square multiplied by the dilution factor and 10^4^, and the total volume (5 mL) of the suspension.

### Metabolite, protein, and lipid extraction (MPLEx)

The extraction procedure was adapted from the Nakayasu et al., 2016 (17). Briefly, the lyophilized fungal biomass resuspended in 100 µL of 18.2 Ω Milli-Q, and 5 volumes of cold (- 20 °C) 2:1 (vol/vol) chloroform-methanol (HPLC grade, Fisher Chemical) solution was added to the samples. Samples were incubated for 5 min on ice, subjected to vortex mixing for 1 min, and centrifuged at 12,000 rpm for 10 min at 4 °C. For metabolomics, the upper aqueous phase, containing hydrophilic metabolites, were collected in glass autosampler vials. To cover a broad spectrum of metabolome, from highly non-polar lipids to extremely polar water-soluble metabolites, 5 µL of the bottom organic phase containing lipids was added to each metabolite sample. The interphase, containing proteins, were washed three times using 500 µL of cold (-20°C) methanol (HPLC grade, Fisher Chemical), vortexed for 1 min, and centrifuged at 10,000 x g for 10 min at 4°C. The supernatants were discarded, and the resulting pellets were dried in a vacuum centrifuge for 5-10 min.

### Sample preparation, LC-MS/MS proteomics data generation and analyses

The dried pellets were dissolved in 100 mM NH4HCO3 containing 8 M urea and the protein concentration was measured by BCA assay (Thermo Scientific cat. no. 23225). Disulfide bonds were reduced by adding dithiothreitol (Thermo Scientific cat. no. A39255) to a final concentration of 10 mM and incubating at 37 °C for 60 min. Samples were then alkylated with a final concentration of 20 mM iodoacetamide (Thermo Scientific, cat. no. A39271) at room temperature (25 °C) for 30 minutes. The reaction was then diluted 10-fold with 100 mM NH_4_HCO_3_ followed by the addition of CaCl_2_ to 1 mM final concentration. Digestion was carried out for 14 hours at 37 °C with 1:50 (wt:wt) trypsin (Promega, cat. no. VA9000) to protein ratio. Salts and reagents were removed by solid-phase extraction using C18 cartridges (Harvard Apparatus, cat. no. 74-4116) according to the manufacturer instructions and the resulting peptides were dried in a vacuum centrifuge.

The LC-MS/MS analysis was performed using a Dionex UltiMate 3000 UHPLC system (Thermo Fisher Scientific, CA, USA) connected to a Q Exactive HF-X mass spectrometer (Thermo Scientific, Bremen, DE). Resuspended tryptic peptides were loaded onto an in-house packed trap column (100 μm I.D., 4 cm long, 5 μm particle size, C18 packing material, Phenomenex) using mobile phase A (0.1% formic acid in HPLC grade water, Fisher Chemical) at a flow rate of 7 μL/min for 5 min and were separated on an analytical column (75 μm I.D., 35 cm long, 1.7 µm particle size, C18 BEH 130 Å packing material, Waters) at 200 nL/min using mobile phase B (0.1% formic acid in HPLC grade acetonitrile, Fisher Chemical). The analytical column was heated to 50 °C and the peptides were separated using a linear gradient from 1% to 7% B over 7.6 min, then 7% to 21% B over 97 min, followed by an increase to 35% B over 8 min. The gradient was subsequently ramped to 75% B in 5 min, then 95% B in 3 min and held at 95% B for 9 min to wash the column. The total LC run time was 190 min including column wash and re-equilibration. The Q Exactive HF-X mass spectrometer was operated as follows: positive polarity, spray voltage of 3.5 kV, funnel RF at 40%, and an inlet capillary temperature of 250 °C. The data-dependent acquisition (DDA) using the full MS-ddMS2 setup was used and the total acquisition time was 120 min. RAW files were recorded using a top 12 DDA method. Full MS scans were acquired with a resolving power of 60,000 with an automatic gain control (AGC) target value of 3E6 charges and maximum injection time (IT) of 20 ms. The mass range was from *m/z* 200 to 2000. The AGC target value for fragmentation spectra was set to 1E5. The isolation width was set to 0.7 *m/z*, and the first mass was fixed at *m/z* 110. Peptide match was set to preferred, and isotope exclusion was on. MS1 and MS2 spectra were acquired in profile mode, respectively. Precursors were fragmented by HCD using 30% normalized collision energy (NCE) and analyzed in the Orbitrap at a resolution of 30,000 with an injection time of 100 ms and an AGC target of 5E4. The dynamic exclusion duration of fragmented precursor ions was set to 45 s. The ion selection abundance threshold was set at 5E3 with charge state exclusion of unassigned and z = 1, and z = 7 to 8 ions.

All raw files were processed by FragPipe (version 22) and searched against the UP000027238 UniProt protein sequence database with decoy sequences (containing 12697 forward entries). Search settings included a precursor mass tolerance of ± 20 ppm, fragment mass tolerance of ± 20 ppm, deisotoping allowing C12/C13 isotope errors of −1/0/1/2/3, strict trypsin and Lys-C as the enzyme (with allowance for N-terminal semi-tryptic peptides), carbamidomethylation as fixed modification, and several variable modifications, including oxidation of methionine, and N-terminal acetylation. Protein and peptide identifications were filtered to a false discovery rate of less than 0.01 within FragPipe. FragPipe result files were then imported into RStudio for downstream analysis in the R environment (version 4.5.1). Median normalization was performed with the “median_normalization” function from the proDA R package (61) across all proteomics datasets. For cases where complete data was needed, imputation was performed with k-nearest neighbors imputation. PCA was performed using PCAtools package (62). Gene set enrichment analysis using the KEGG pathway database was performed with the clusterProfiler R (gseKEGG function) (63, 64) package using *Colletotrichum higginsianum* reference (UP000092177) annotations, after BLASTP annotation transfer, while gaussian mixture modeling (GMM) clustering was accomplished with the Mclust R (65) package. Venn diagrams were generated using eulerr (66), and UpSet plots were generated using UpSetR (67).

### LC-MS/MS metabolomics data generation and analyses

Metabolite extract (ME) was analyzed using LC-MS/MS using a Vanquish Flex UHPLC system (Thermo Scientific, interfaced with a Q Exactive HF-X (Thermo Scientific, Bremen, DE) mass spectrometer. For reversed-phase (RP) C18 liquid chromatography-tandem mass spectrometry, same volume (5-10 µL) of ME are injected and compounds are separated using a Hypersil GOLD column (2.1 mm I.D., 150 mm long, 3 μm particle size, Thermo Scientific) with a column temperature of 40 °C. Mobile phase A (0.1% formic acid in HPLC grade water, Fisher Chemical) and mobile phase B (0.1% formic acid in HPLC grade acetonitrile, Fisher Chemical) were initially 90% and 10%, respectively, with a flow rate of 400 μL min^−1^. The gradient method continued as follows: 0 to 2 min at 90% A with a flow rate of 400 μL min^−1^; 2 to 11 min decreased to 10% A with a flow rate of 400 μL min^−1^; 11 to 12 min held at 10% A with a flow rate of 400 μL min^−1^; 12 to 12.5 min held at 10% A with an increased flow rate to 500 μL min^−1^; 12.5 to 13.5 min increased to 90% A with a flow rate of 500 μL min^−1^; 13.5 to 14 min held at 90% A with a flow rate of 500 μL min^−1^; 14 to 14.5 min held at 90% A with a decreased flow rate to 400 μL min^−1^; and 14.5 to 16 min held at 90% A with a flow rate to 400 μL min^−1^. For hydrophilic interaction liquid chromatography (HILIC) tandem mass spectrometry chromatography was adapted from Clendinen et al., 2019 (63), and 5 to 10 µL of ME are injected and compounds are separated using a using ACQUITY UPLC BEH HILIC column (2.1 mm I.D., 100 mm long, 1.7 μm particle size, Waters) with a column temperature of 50 °C. Mobile phase A (5% acetonitrile: 95% 10 mM NH_4_OAc in water with 0.05% NH_4_OH) and B (100% acetonitrile with 0.05% NH_4_OH) were initially 5% and 95%, respectively, at a flow rate of 300 μL min^−1^. The gradient method continued as follows: 0 to 6.0 min at 63% A with a flow rate of 300 μL min^−1^; 6.0 to 7.0 min held at 63% A with a flow rate of 300 μL min^−1^; 7.0 to 7.1 min decreased to 5% A with a flow rate of 300 μL min^−1^; 7.1 to 7.2 min held at 5% A with an increased flow rate to 500 μL min^−1^; 7.2 to 9.5 min held at 5% A with a flow rate of 500 μL min^−1^; 9.5 to 9.7 min held at 5% A with a decreased flow rate of 300 μL min^−1^; and 9.7 to 12.0 min held at 5% A with a flow rate of 300 μL min^−1^.

For both separations the samples were analyzed in both positive and negative ionization modes using HCD. The HESI source parameters are set as follows: spray voltage of 3.7 kV or 3.0 kV for positive and negative modes, respectively, funnel RF at 50%, an inlet capillary temperature of 350 °C, and auxiliary gas heater temperature set to 150 °C. Full MS scans were acquired with a resolving power of 120,000 from *m/z* 80 to 800, an AGC target of 3E6 ions, with a maximum injection time of 20 ms. The DDA (dd-MS2) parameters used to obtain product ion spectra are as follows: resolving power was set to 15,000, an AGC target of 1E5 ions, with maximum injection time of 100 ms, isolation width of 0.4 *m/z*, loop count 12, and NCE set to 20, 40, and 60 eV. For data generated, data was processed in Compound Discoverer 3.3 (Thermo Scientific) to confidently identify metabolites. For positive and negative mode RP and HILIC datasets, spectra are aligned using an adaptive curve with a maximum of 0.3 or 0.6 RT shift respectively and a 3 ppm mass tolerance. Peaks are selected based on a minimum intensity of 1.5E5 and a chromatographic S/N of 3. Detected features are grouped based on a mass tolerance of 3 ppm and a RT tolerance of 0.3. Features were then filtered out if not present in at least 75% samples. Compounds were identified via on internal and external database matches and assigned based on Isotopic pattern, RT (internal database only), MS1, and/or MS2. All identifications and integrated peaks were manually validated and exported for statistical analysis. The downstream analysis was performed using MetaboAnalyst 6.0 (64) and in R version 4.5.1.

## Funding

This work by the DOE Office of Science Biological and Environmental Research (BER), under KP1601011 LAB-23-2955 and performed on project award (61073, L.P.-T.) from EMSL, a DOE Office of Science User Facility sponsored by the Biological and Environmental Research program under Contract No. DE-AC05-76RL01830.

## Acknowledgements

We thank Dr. Louis K. Prom from the USDA-ARS and Dr. Clint W. Magill of Texas A&M University for providing *C. sublineola* pathotype *FSP237*. We also thank Dr. Matthew E. Monroe of Pacific Northwest National Laboratory for his assistance in depositing the raw proteomic data onto MassIVE.

## Data Availability Statement

The mass spectrometry raw data and database search results are accessible through the ProteomeXchange Consortium and MassIVE data repository with dataset identifiers MSV000101534. Associated code for the data analysis, result tables, metadata used for figure generation can be found at the GitHub repository (https://github.com/prdawar/Cs_nutrient_limitation_stress).

## Conflict of Interest

Authors have no conflict of interest.

## Author contributions

Pranav Dawar -Conceptualization, Methodology, Formal analysis, Investigation, Visualization, Writing *-*original draft, Writing *-*review & editing

Dora Farago -Methodology, Writing *-*review & editing

Kevin J. Zemaitis -Conceptualization, Methodology, Writing *-*review & editing

Audrey Thomas -Methodology, Writing *-*review & editing

Priscila M. Lalli -Methodology, Writing *-*review & editing

Chaevien S. Clendinen -Methodology, Formal analysis, Writing *-*review & editing

Vanessa L. Paurus -Methodology, Writing *-*review & editing

Theresa F. Law -Methodology, Writing *-*review & editing

Erin L. Bredeweg -Conceptualization, Methodology, Writing *-*review & editing

James M. Fulcher -Conceptualization, Methodology, Writing *-*review & editing

Jeffery L. Dangl -Conceptualization, Writing *-*review & editing

Qun Liu -Conceptualization, Writing *-*review & editing

Ljiljana Paša-Tolić - Conceptualization, Funding acquisition, Methodology, Project administration, Resources, Supervision, Writing *-*review & editing

## Supplementary Information

**Supplementary Figure 1 -** Sample-sample correlation heatmap of protein abundance profiles. Pearson correlation coefficients were calculated between all samples using the QC-verified proteins, with missing values handled using pairwise complete observations. Correlations are displayed as a shaded correlogram, where darker blue indicates stronger positive correlation. The diagonal represents self-correlation for each sample. Red rectangles highlight predefined sample groupings in the matrix.

**Supplementary Figure 2 -** A) Density histograms showing the distribution of log2-transformed Cluster 18 protein intensities across conditions. B) Volcano plot of Cluster 18 proteins across comparisons. Dashed lines mark |log2 fold-change| = 1 and FDR = 0.05. C) Density histograms showing the distribution of log2-transformed Cluster 13 protein intensities across conditions. D) Volcano plot of Cluster 13 proteins across comparisons. Dashed lines mark |log2 fold-change| = 1 and FDR = 0.05.

**Supplementary Data S1 -** Conidia counts per condition. Bioreps (n) = 5

**Supplementary Data S2 -** One-tenth-strength PDA specific proteins identified in *Cs*

**Supplementary Data S3 -** List of significantly differentially abundant proteins identified from one-tenth-strength PDA and half-strength PDA condition comparison, using log2FC ≥ 1 and adjusted p < 0.05 cut-offs.

**Supplementary Data S4 -** List of significantly differentially abundant proteins identified from one-tenth-strength PDA and full-strength PDA condition comparison, using log2FC ≥ 1 and adjusted p < 0.05 cut-offs.

**Supplementary Data S5 -** List of significantly differentially abundant proteins identified from half-strength PDA and full-strength PDA condition comparison, using log2FC ≥ 1 and adjusted p < 0.05 cut-offs.

**Supplementary Data S6 -** Complete list of proteins mapped to Cluster 8. Column E-H represent the significant DAPs, using log2FC ≥ 1 and adjusted p < 0.05 cut-offs.

**Supplementary Data S7 -** List of significantly differentially abundant metabolites identified from half-strength PDA and full-strength PDA condition comparison, using log2FC ≥ 1 and adjusted p < 0.05 cut-offs.

**Supplementary Data S8 -** List of significantly differentially abundant metabolites identified from one-tenth-strength PDA and full-strength PDA condition comparison, using log2FC ≥ 1 and adjusted p < 0.05 cut-offs.

**Supplementary Data S9 -** List of significantly differentially abundant metabolites identified from one-tenth-strength PDA and half-strength PDA condition comparison, using log2FC ≥ 1 and adjusted p < 0.05 cut-offs.

## Notes

### Competing Interest Statement

The authors have declared no competing interest.

### Summary of Updates

This version of the manuscript has been revised to improve the following - 1) Corrected the arrow colors in Figure 3. 2) Corrected the graphical abstract. 3) Improved the overall quality of the main text figures.

