## Supplementary Figure 1 and 2 for "Studying Effects of PDA Media Strengths on the Growth of *Colletotrichum sublineola* Using MPLEx-Based Integrative Proteomics and Metabolomics Analyses"

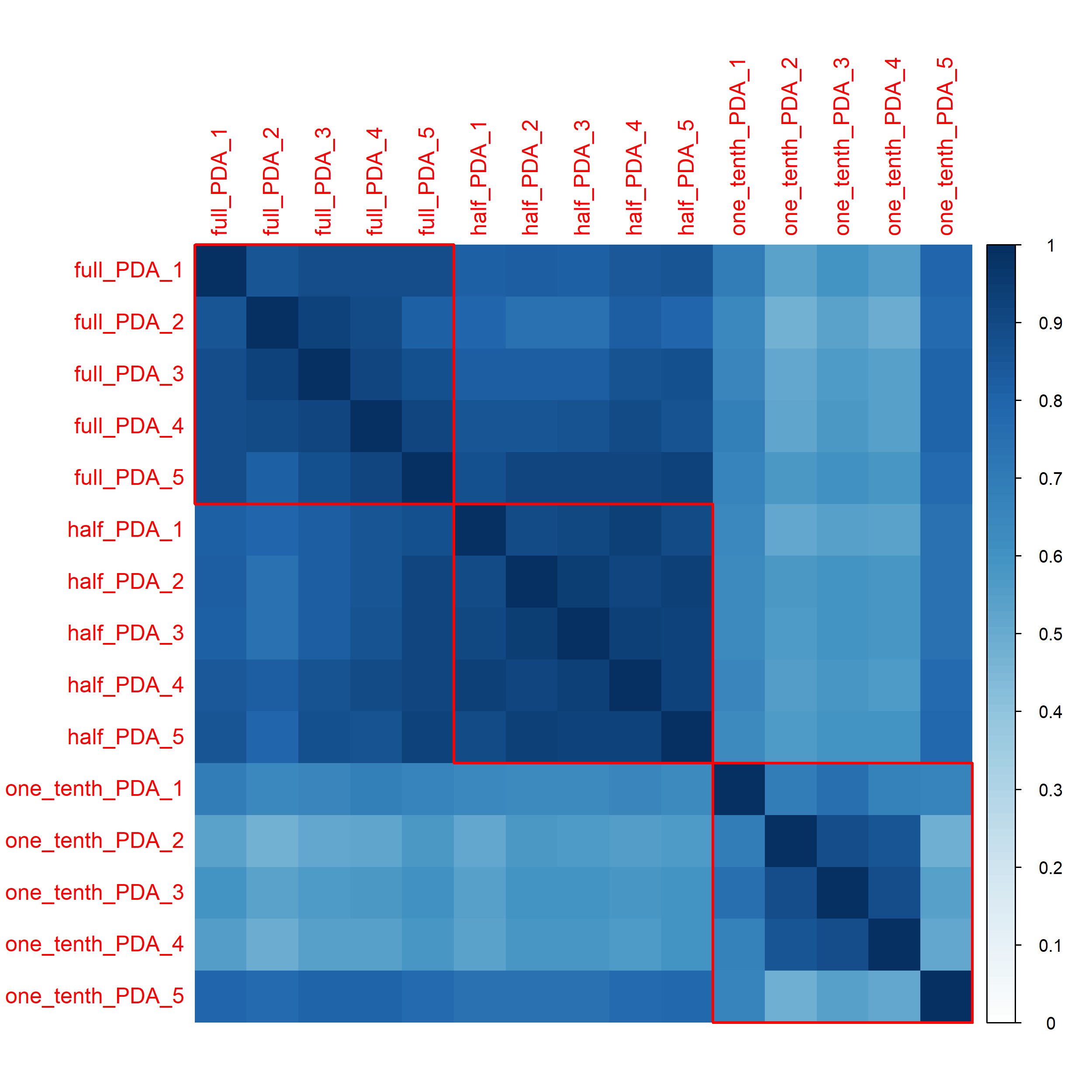

**Supplementary Figure 2**

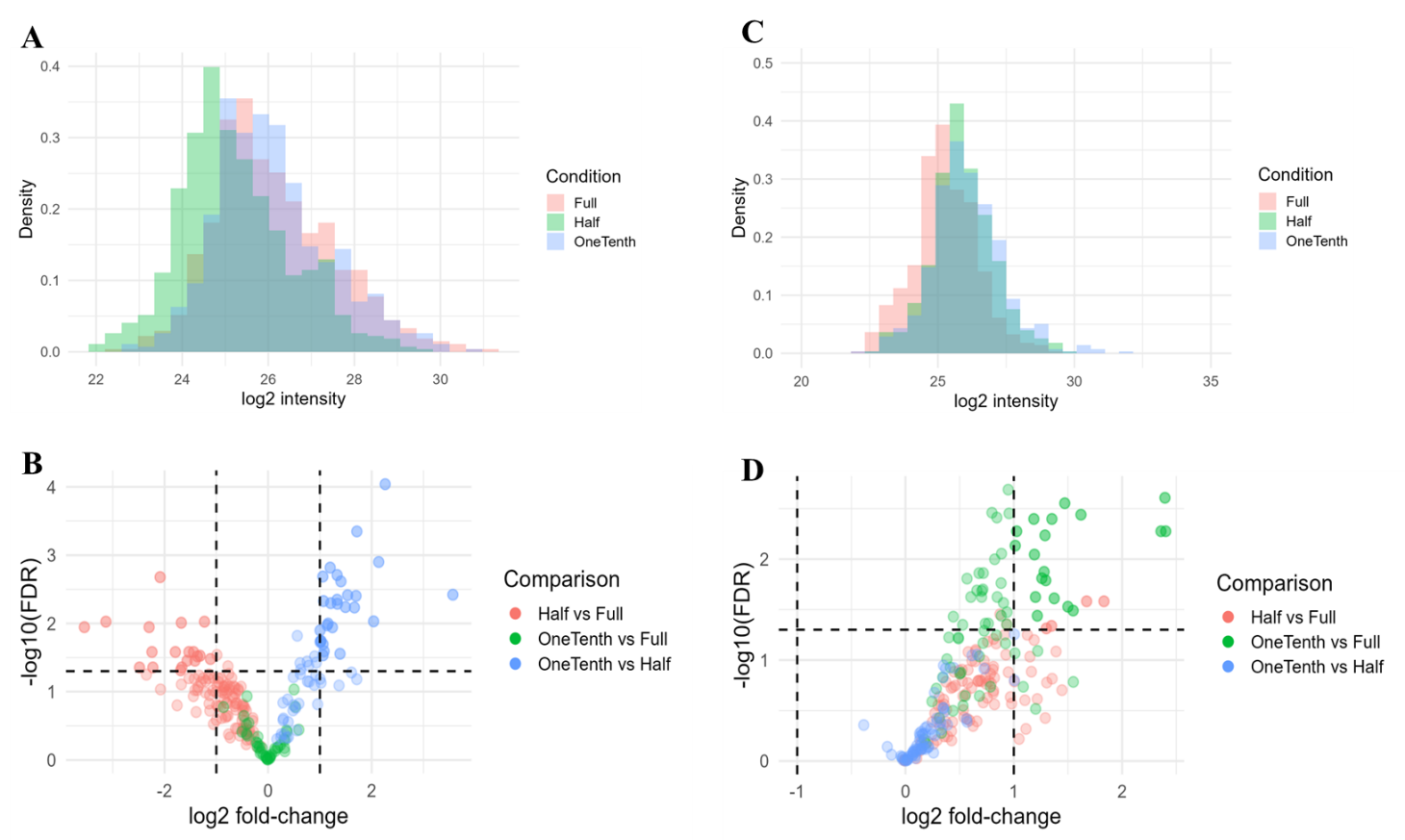
